# A Generalizable Deep Learning Model for Automated 3D Segmentation of Orthopteran Head Anatomy in Micro-CT

**DOI:** 10.64898/2026.08.12.744546

**Authors:** Arthur Chéron, Shinichi Morita, Naoki Morimoto, Takahiro Ohde

**Affiliations:** Department of Applied Biosciences, Graduate School of Agriculture, Kyoto University, Japan; Division of Evolutionary Developmental Biology, National Institute for Basic Biology, Japan; Department of Basic Biology, School of Life Science, The Graduate University for Advanced Studies, SOKENDAI, Japan; Laboratory of Physical Anthropology, Graduate School of Science, Kyoto University, Japan

**Keywords:** Orthoptera, automated segmentation, micro-CT, deep learning, nnU-Net, comparative biology, *Gryllus bimaculatus*, *Loxoblemmus equestris*, *Loxoblemmus doenitzi*, UNet

## Abstract

Deep learning tools are increasingly used today, particularly in medical segmentation. A gap nonetheless remains in automating segmentation for insects. This work addresses the following question: can a generalist segmentation model, trained on several phylogenetically related orthopteran species, reliably automate head tissue segmentation from micro-CT images? To answer this, we used nnU-Net, a self-configuring 3D deep learning segmentation framework originally developed for medical imaging, whose core function, learning to recognize tissues of interest, applies directly to this context. Six anatomical classes were automated, comparing two training strategies: sequential fine-tuning, which adds species one at a time under the assumption that progressive learning would strengthen predictive power, and from-scratch training, in which the model learns the entire dataset simultaneously. The fine-tuning model (ModelB) reached a Dice coefficient (a measure of overlap between automated segmentation and manual ground truth, ranging from 0 to 1) of 0.7715, compared to 0.7664 for the from-scratch model (ModelC). Although both models produced accurate automated segmentations, no significant difference was found between the two training strategies (paired Wilcoxon test, n = 24, p = 0.243). Despite a dataset limited to 20 individuals and the absence of one method clearly outperforming the other, the models remain usable across the three species studied (*Gryllus bimaculatus*, *Loxoblemmus equestris*, *L. doenitzi*), including in the presence of pronounced sexual dimorphism. It reduces a 20 hour segmentation task to under a minute.

## 1 Introduction

Anatomical tissue segmentation is a technique widely used in research, whether applied to insects, plants, fossils, or, most commonly, medical imaging. This method allows researchers to visualize and quantitatively analyze regions of interest without dissection. The main obstacle to 3D modeling is the time required for manual segmentation of images (micro-CT, MRI, CT scan, and so on). Segmenting every tissue in the head of a single specimen can take more than twenty hours, which represents a real bottleneck for researchers, demanding labor, anatomical expertise, and time. There is also a significant bias to anticipate in manual segmentation due to the possibility of repeatedly missing certain tissues.

Given the strong demand from the medical field, many models have been proposed for the automated segmentation of organs or tumors. What all of these models have in common is their reliance on nnU-Net, a self-configuring 3D deep learning segmentation framework that has made this kind of automation considerably more accessible [1]. Building a comparable system for Orthoptera is a genuine challenge worth pursuing, given their beneficial and harmful effects on crops, their nutritional value, and their importance for understanding insect developmental biology. Mole crickets, for instance, illustrate this well: their extensive digging of vertical and horizontal galleries reshapes soil structure, increasing water infiltration while also accelerating soil erosion [2]. In addition, the domestication of cricket species, combined with a warming climate, could represent a new phytosanitary threat to seed crops [3]. This dual role, alternating between soil engineer and potential pest, highlights the importance of better understanding the fundamental biology of Orthoptera.

This is why developing a model capable of automating the segmentation of Orthoptera heads based on a deep learning method developed for medical imaging could fill this existing gap. A similar approach was used in the work of Toulkeridou et al. [4] on numerous ant species. However, their model consists of a single class of interest (the brain) using the 2D U-Net architecture, thereby limiting spatial context and producing predictions slice by slice rather than as an entire volume.

This raises a question that goes beyond a simple technical issue: can a generalist segmentation model be trained across several orthopteran species? This is a fundamental question for comparative computational biology, since the answer determines whether segmentation pipelines can realistically scale to the breadth of biological diversity. Toulkeridou et al. [4] demonstrated, in a quantitatively rigorous way, that a brain segmentation model could generalize to ant species absent from training (up to 80% IoU, 90% Dice across 30 test species), but remained confined to a single anatomical structure.

In this study, we build a 3D, multi-class automated segmentation model for the orthopteran head in micro-CT, capable of simultaneously segmenting six anatomical structures: muscles, esophagus, brain, eyes, external cuticle, and internal cuticle. To achieve this, we developed and evaluated an iterative fine-tuning strategy applied group by group across species, in which a base model trained on *G. bimaculatus* is progressively adapted to *L. equestris*, then *L. doenitzi*, with the possibility of extending it further to other species in the future.

## 2 Materials and Methods

### 2.1 Specimens and sample preparation

*Gryllus bimaculatus*, *Loxoblemmus equestris*, and *Loxoblemmus doenitzi* specimens were all reared at the Laboratory of Insect Physiology, Kyoto University, under standard controlled conditions (a constant 29°C day and night, a 16h/8h light cycle, and a diet of Tetra brand goldfish flakes).

Head preparation followed the same protocol as Yoneda et al. [5], with one modification: staining with 1% Lugol’s solution (I_2_) for 48 hours.

### 2.2 X-ray micro computed tomography (micro-CT) acquisition

Scans were acquired at voxel resolutions ranging from 0.0015 to 0.0044 mm depending on the specimen (each voxel corresponds to a cube measuring 1.5 to 4.4 micrometers on a side), producing reconstructed volumes ranging from 278x312x257 to 1008x1008x630 voxels depending on the size of the scanned individual. Reconstructed volumes were saved in NRRD format, a standard format for volumetric medical images. Two scanners were used: a VoxelWorks VWHTA-23003 (80 kV, 100 *µ*A) at Kyoto University, and a Bruker SKYSCAN 1272 (50 kV, 80 *µ*A) at the National Institute for Basic Biology (NIBB).

### 2.3 Manual reference segmentation

Segmentation consists of assigning each voxel in the volume a label corresponding to the anatomical structure it belongs to (muscle, brain, and so on), or, for everything else, a generic label for anything that is not a structure of interest, named *Others/ Background*. It is this manual segmentation that serves as ground truth for training and evaluating the automated model.

Segmentation was performed in 3D Slicer (version 5.10.0), an open-source platform for visualizing and annotating volumetric medical images [6]. The method used was *Grow from Seeds*: the user paints representative strokes for each structure every few slices (the exact interval depending on the total number of slices in the image set), and the algorithm propagates these annotations across the rest of the volume based on the similarity of color intensity and texture between voxels. This intermediate automated prediction was then corrected manually, slice by slice, across the three orthogonal viewing planes (axial, coronal, sagittal), and finally checked in 3D, to obtain a precise segmentation. To carry out assisted segmentation, a Python script (sparse_seg.py, see appendix) was applied to the segmentation produced by one of the models, converting the 3D volume into a set of slices (chosen according to the image set) along one or several axes. Each resulting slice could then be corrected individually before applying the Grow from Seeds tool again in 3D Slicer. Only the training specimens benefited from this assisted segmentation, in order to avoid biasing Dice coefficients through a model-driven similarity effect on the validation and test specimens.

Six anatomical structures of interest were annotated: muscles, esophagus, brain (including the optic lobes, ocellar lobes, and nerves), eyes, external cuticle, and internal cuticle. Label 0 groups together all unannotated voxels (*Others/Background*).

### 2.4 Data preprocessing and formatting

Manual segmentation files and their corresponding raw images were converted from NRRD format to NIfTI format (.nii.gz extension, via the convert_specimen.py script, see appendix), the input format required by the machine learning software used. This conversion was automated using Python scripts developed specifically for this project, relying on the SimpleITK library. This step included two main operations: (i) harmonizing labels across specimens according to the shared scheme described above; (ii) resampling the raw image so that it matched exactly the geometric space of its segmentation, using linear interpolation for the grayscale image.

### 2.5 Deep learning model principle and configuration

Deep learning refers to a family of artificial intelligence methods built on multi-layer artificial neural networks, capable of learning to recognize complex patterns directly from example data, without explicitly defining recognition rules.

For image segmentation, the reference architecture is U-Net, a type of neural network originally developed for medical image segmentation [7]. It relies on two complementary pathways: a contracting path (encoder) that progressively reduces image resolution while increasing the number of abstract features extracted (allowing the network to grasp global context), followed by a symmetric expanding path (decoder) that progressively reconstructs a full resolution output image, in which each voxel is assigned a predicted class (muscle, esophagus, and so on).

This project specifically uses nnU-Net (v2.7.0), a system that automates the configuration of the U-Net network [1]. Unlike a manual U-Net setup, where the user has to choose numerous technical parameters (input image size, number of layers, learning rate, and so on), nnU-Net automatically analyzes the characteristics of the dataset provided (image size, voxel spacing, number of classes) and configures a suitable architecture on its own. This tool makes deep learning particularly accessible to users.

In this work, the configuration chosen is the fully three dimensional version of the network (3D fullres), which processes 3D volumes directly rather than individual 2D images, unlike the approach used by Toulkeridou et al. [4], which processed individual 2D slices before reconstructing the final volume. Working directly in three dimensions allows the network to exploit the full spatial context (the position of a class relative to its neighbors in all three dimensions).

The configured architecture has six encoding stages, with the number of channels (features extracted at each stage) growing from 32 to 320. Each prediction is made on a patch (portion) of the volume measuring 112 *×* 128 *×* 160 voxels, with 2 patches processed simultaneously at each training step (batch size).

Model training was carried out on a workstation equipped with an Intel Core Ultra 7 255HX processor, an NVIDIA GeForce RTX 5070 GPU, 12 GB of VRAM, and 32 GB of RAM, running PyTorch 2.11.0 (CUDA 12.8). The patch size (112 *×* 128 *×* 160 voxels) and batch size were determined automatically by nnU-Net based on the available GPU memory.

Training consists of iteratively adjusting the network’s internal parameters to minimize a loss function measuring the gap between the network’s predictions and the manually annotated ground truth. This loss function combines Dice loss (derived from the Dice coefficient, see Section 2.7) and cross entropy (a classic statistical measure of the quality of a classification). Parameter adjustment is carried out using the Adam optimizer, with a learning rate (the magnitude of the adjustments made at each step) starting at 0.01 and decreasing progressively over the course of training, which consists of 1000 passes (epochs).

### 2.6 Iterative multi-species fine-tuning strategy

A straightforward approach would have been to train a single model, in one pass, on all available specimens across every species at once (the *from-scratch* method). An alternative strategy, known as fine-tuning, was favored in this work.

The principle behind fine-tuning is to start from a model that has already been trained and adapt it progressively to new data, rather than starting from zero. This approach rests on the idea that the parameters learned to recognize the general anatomy of a cricket remain largely relevant for a related species, requiring only light adjustment rather than being relearned entirely.

Over the course of this research, an initial series of models was developed iteratively (ModelA), progressively adding specimens as they were annotated. This exploratory approach was mainly useful for generating preliminary predictions that facilitated the assisted annotation of new specimens. The formal results presented below concern two controlled and directly comparable training strategies (ModelB and ModelC), built on the final, complete dataset.

### 2.7 Validation and quantitative evaluation

To allow a consistent comparison across every successive version of the model, one specimen per species was systematically excluded from training at each stage and used only to evaluate performance (the validation and test set). This choice makes it possible to track performance changes on these same specimens across the different versions of the model, without bias linked to a change of specimen.

Model performance was quantified using two standard segmentation metrics, computed separately for each anatomical structure:

The Dice coefficient (*Dice Similarity Coefficient*, DSC) measures the overlap between the model’s prediction and the manual ground truth:

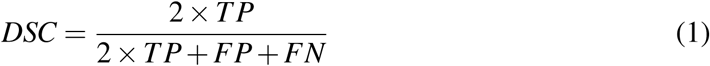

The Jaccard index (IoU) measures the ratio between the region where the prediction and the ground truth overlap and the total area covered by either:

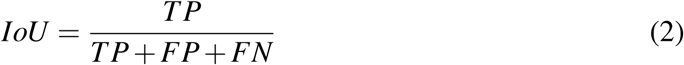

where TP (true positives) refers to voxels correctly identified as belonging to a given structure, FP (false positives) refers to voxels wrongly assigned to that structure, and FN (false negatives) refers to voxels of that structure missed by the model. Both metrics range from 0 (no overlap) to 1 (perfect overlap).

### 2.8 Computational tools

Many tools were used over the course of this work, in particular 3D Slicer [6], which was used for assembling the image datasets, segmenting the anatomical classes, and running the Grow from Seeds tool to produce a precise manual segmentation. RStudio was used for the statistical calculations in R and for generating Figure 5. Anaconda was also used as a Python environment manager. All the Python scripts used for format conversion, label remapping, and metric calculation were developed with the assistance of Claude, Anthropic’s conversational artificial intelligence, used as a resource providing programming knowledge and technical troubleshooting throughout this project.

## 3 Results

### 3.1 ModelB performance

ModelB reaches an overall mean Dice coefficient of 0.7715, as shown in Figure 3. InCuticle is the weakest class, ranging from 0.3019 to 0.5744 across specimens, with an average of 0.4732. Eyes, on the other hand, are the strongest performing class, ranging from 0.9074 to 0.9564, averaging 0.9365. More broadly, the classes Eyes, Brain, and Muscles show the highest Dice coefficients across the model. The first specimen to be segmented (*G. bimaculatus* 01) has the lowest mean Dice score among the four validation specimens (0.7557).

**Figure 1:**
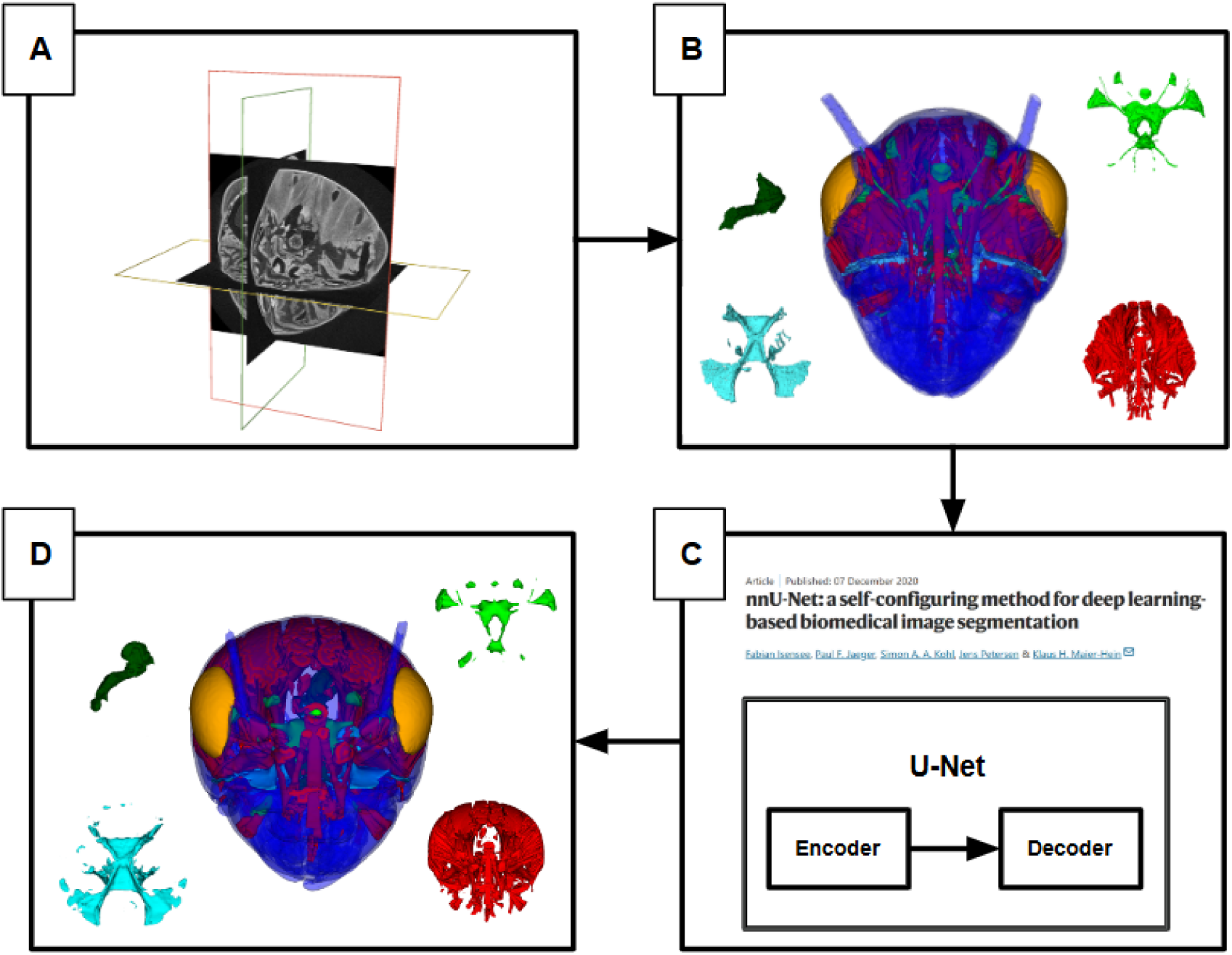
Simplified overview of the model training method. (A) Assembly of CT images in 3D Slicer. (B) Manual or semi-manual segmentation. (C) Data transfer to nnU-Net for training (see Section 2.5). (D) Automated segmentation produced by nnU-Net.

**Figure 2:**
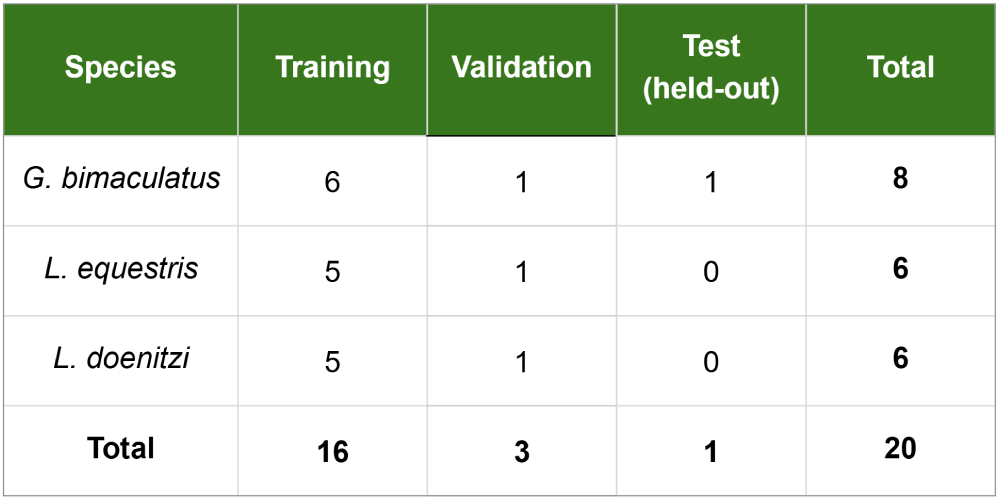
Summary table of the individuals used to build the model (the full table is available in the appendix).

**Figure 3:**
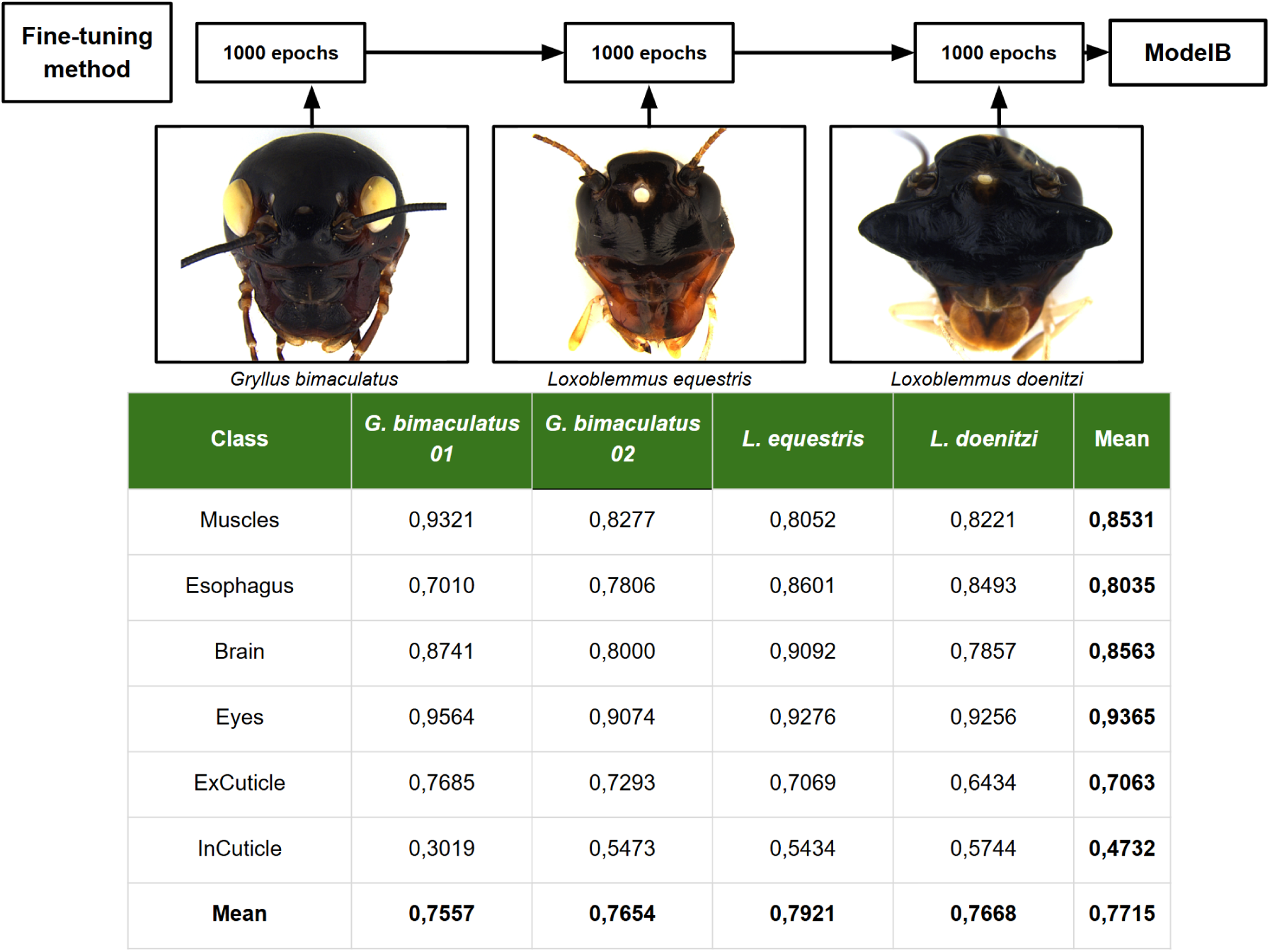
Schematic overview and Dice scores obtained for ModelB (fine-tuning method).

### 3.2 ModelC performance

ModelC reaches an overall mean Dice coefficient of 0.7664, as shown in Figure 4. InCuticle is again the weakest class, ranging from 0.2808 to 0.5852, averaging 0.4635. Eyes remain the strongest class, ranging from 0.9123 to 0.9571, averaging 0.9374. The first specimen segmented (*G. bimaculatus* 01) again shows the lowest mean Dice score among the four validation specimens (0.7426).

**Figure 4:**
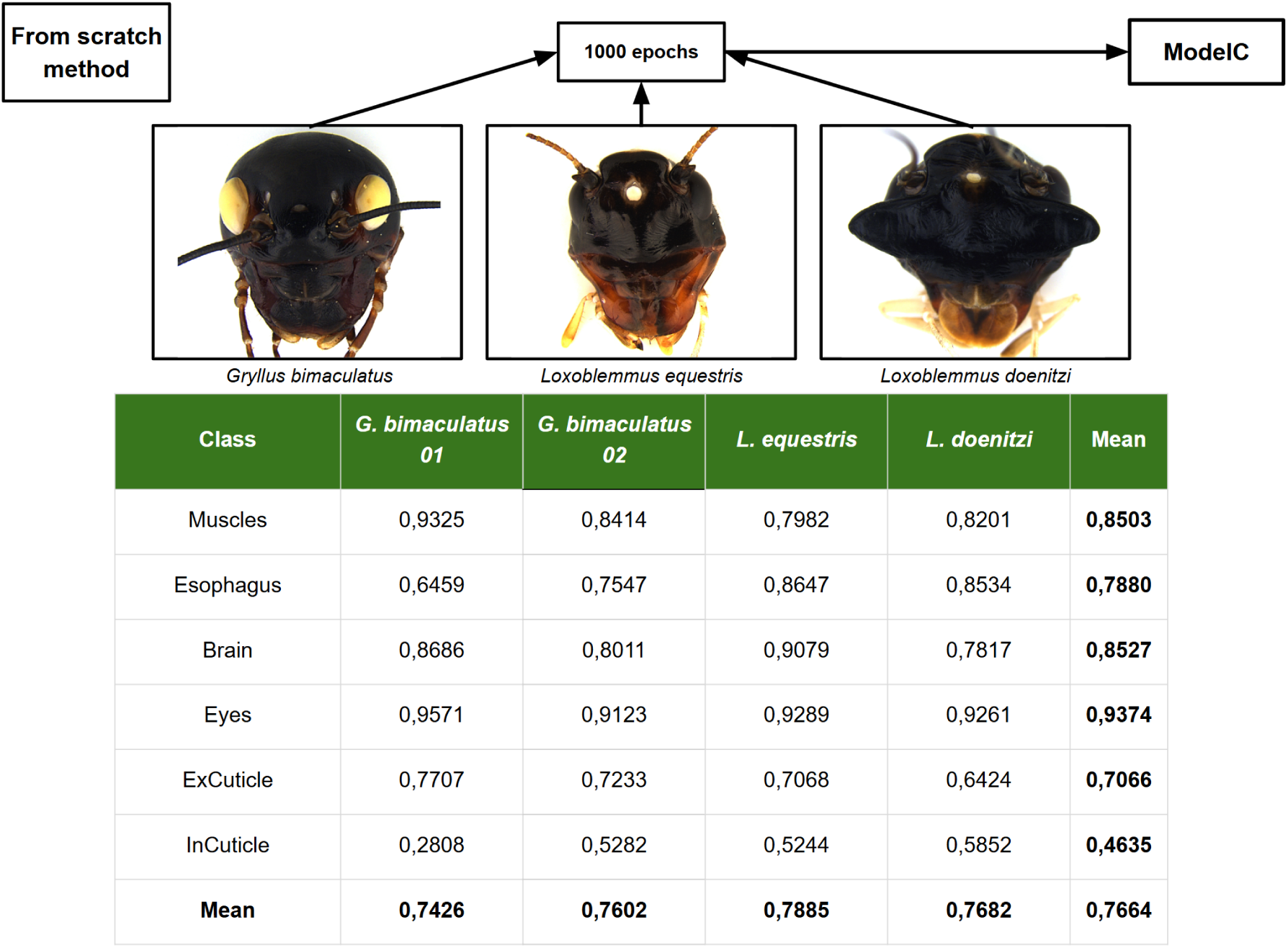
Schematic overview and Dice scores obtained for ModelC (from scratch method).

**Figure 5:**
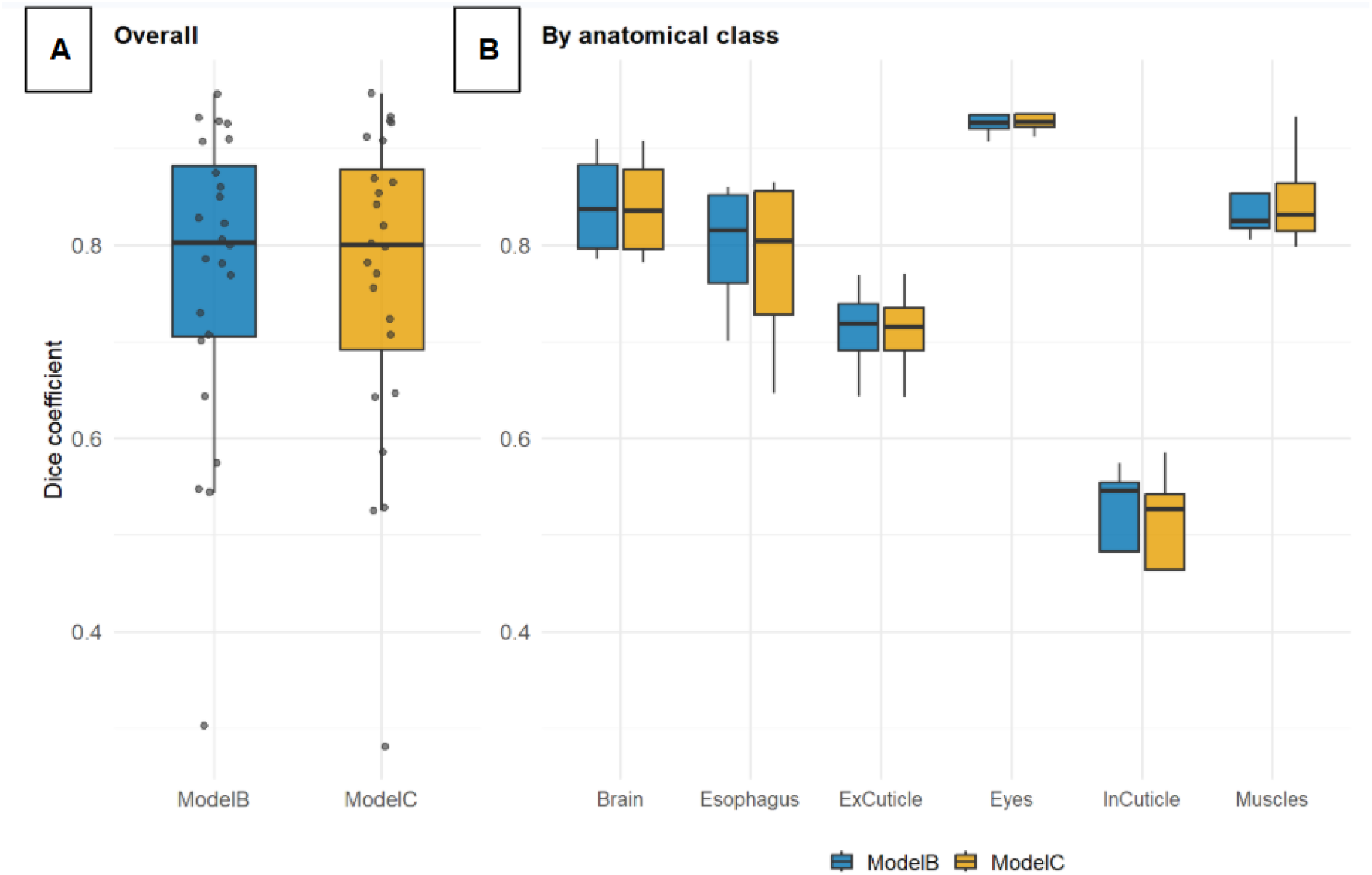
Boxplots comparing Dice coefficients between ModelB (fine-tuning) and ModelC (*from scratch*) (A), and across anatomical classes (B). No significant difference was found (paired Wilcoxon test, n = 24, V = 191.5, p = 0.243).

### 3.3 Comparison of the different training strategies

To determine whether the order in which species are introduced during training affects final model performance, Dice scores from ModelB and ModelC were compared across 24 matched observations (6 anatomical classes *×* 4 validation specimens, including a second *G. bimaculatus* specimen added to strengthen statistical power).

A paired Wilcoxon signed-rank test revealed no statistically significant difference between the two strategies (V = 191.5, p = 0.243; mean difference = *−*0.005). As shown in Figure 5, Dice scores were nearly identical across the two models for the majority of class-specimen combinations.

Similarly to the Dice coefficients obtained, the IoU of ModelB (0.6487) is close to that of ModelC (0.6436), as shown in Figure 6. Following the same ratio, InCuticle remains the weakest class regardless of the metric considered (ModelB = 0.3326, ModelC = 0.3228).

**Figure 6:**
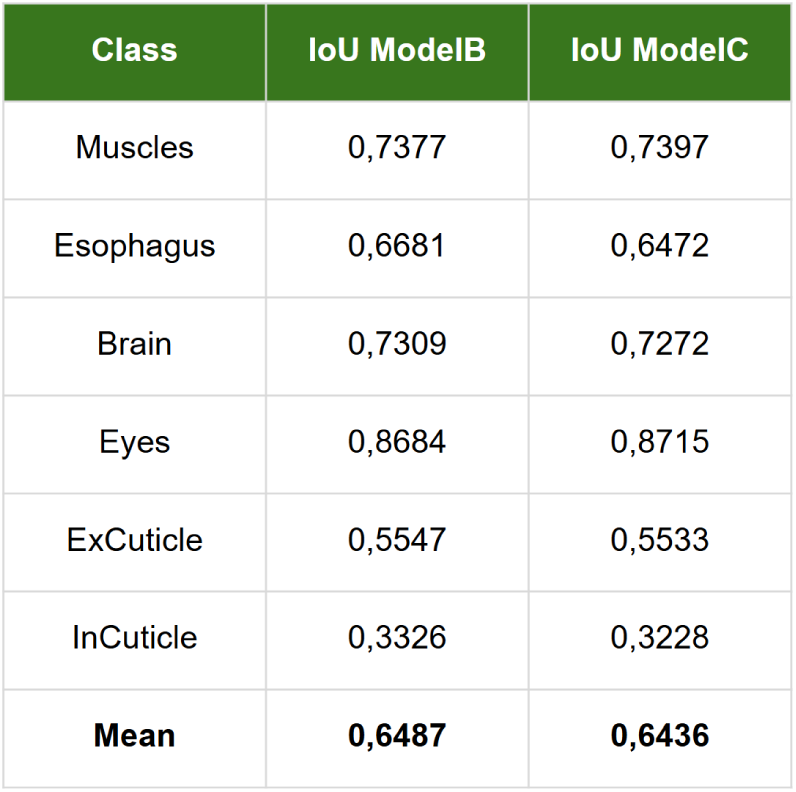
Mean Jaccard index (IoU) by class, ModelB vs ModelC (derived from each individual Dice value via *IoU* = *Dice/*(2 *−Dice*), then averaged).

### 3.4 Generalization beyond Orthoptera

As shown in Figure 7, ModelB (trained exclusively on head anatomy) produces a moderately accurate segmentation of the cuticle and muscles across the entire body of the ant, while the remaining classes remain both unusable and impossible to interpret visually.

**Figure 7:**
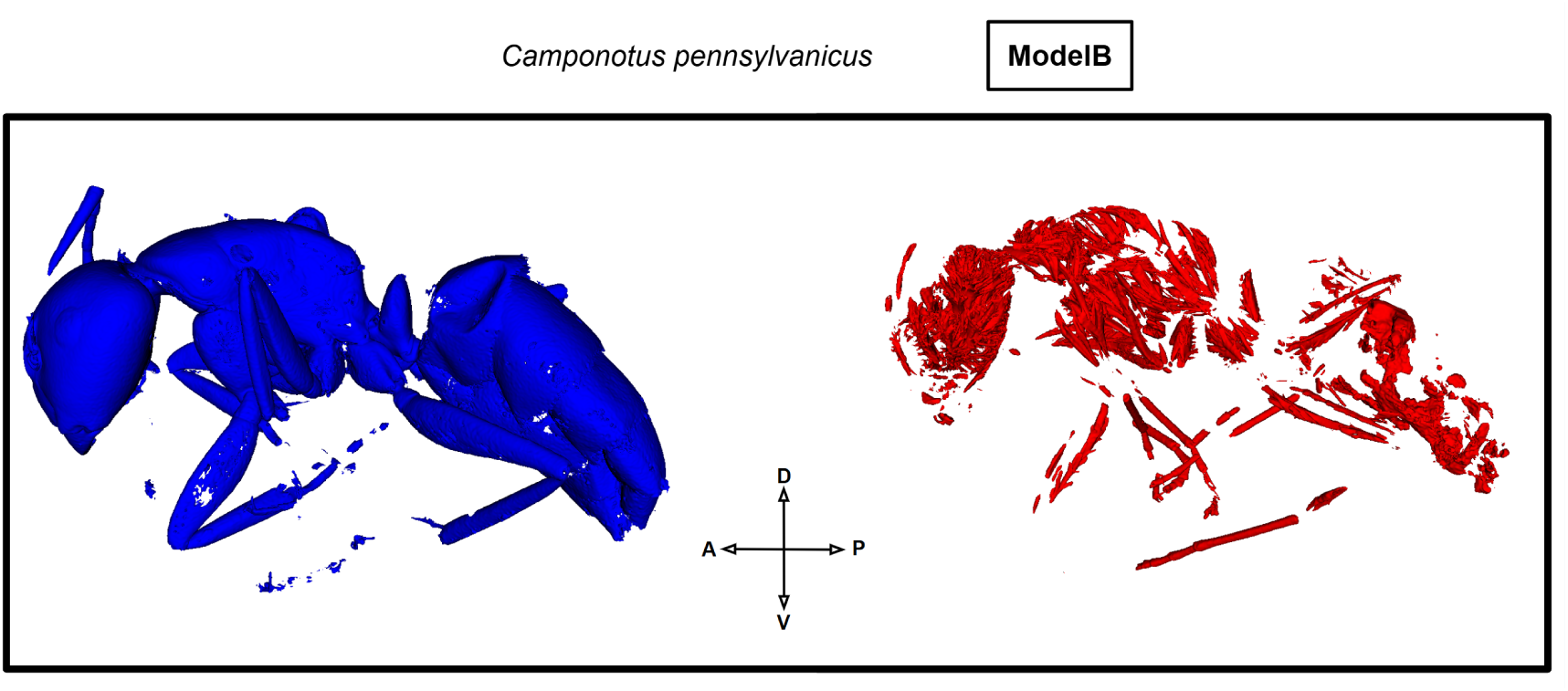
Generalization test of ModelB on a *Camponotus pennsylvanicus* worker ant (automated segmentation: external cuticle in blue, muscles in red).

### 3.5 Exploratory extension to thoracic muscle segmentation

As an exploratory extension beyond head anatomy, the fine-tuning approach was further applied to the segmentation of thoracic muscle groups in *G. bimaculatus*, comprising two segmentation classes of interest: the prothoracic muscles (first thoracic segment) and the meso/metathoracic muscles (the last two thoracic segments), as shown in Figure 8. Three pretraining conditions were compared: a model pretrained via ModelA on two species (t1: *G. bimaculatus* and *L. equestris*), a model pretrained via ModelC on three species (t2: *G. bimaculatus*, *L. equestris*, and *L. doenitzi*), and a model trained from scratch. Mean Dice and IoU scores are highest for t1, while the from-scratch training yields the lowest means.

**Figure 8:**
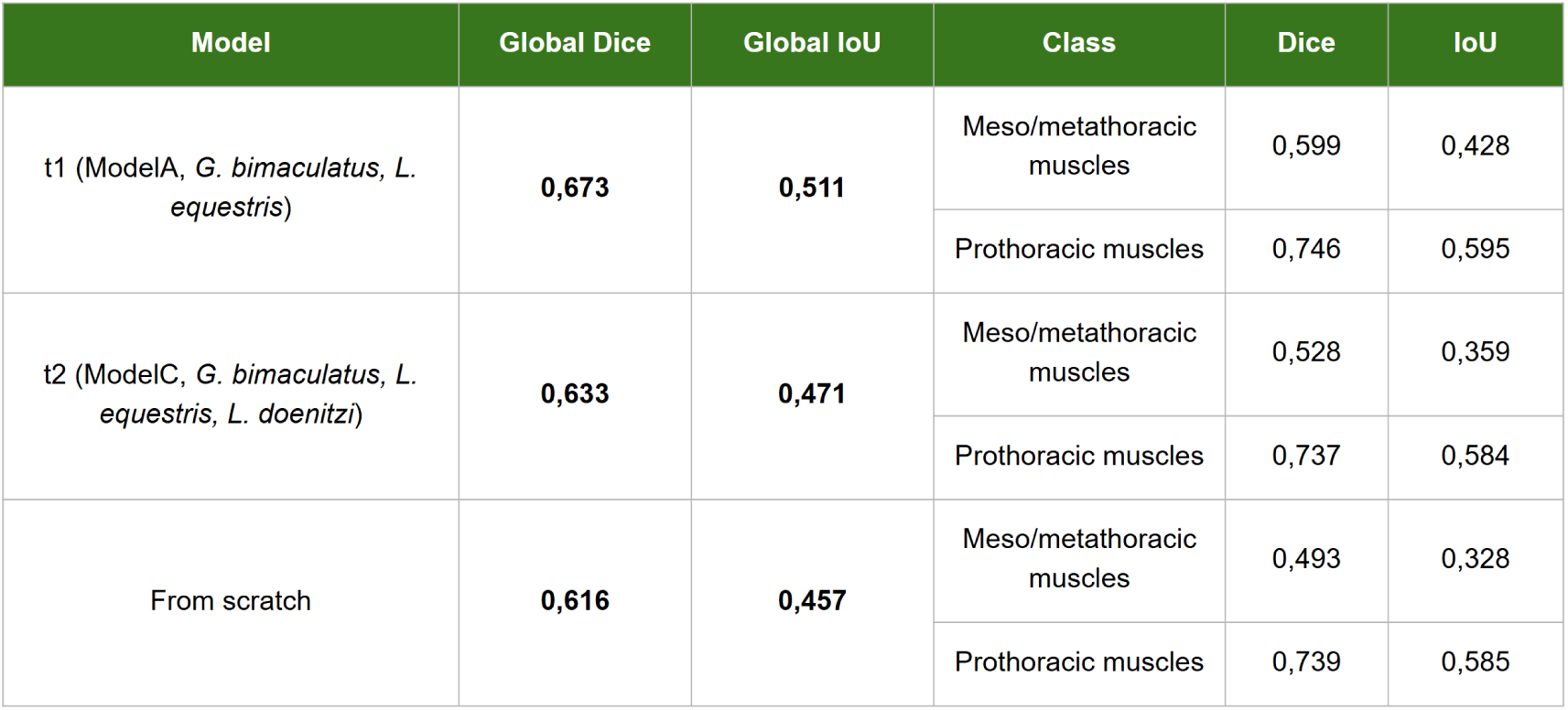
Dice and IoU scores obtained on the thoracic muscle segmentation task for three pretraining conditions: t1 (pretrained on two species), t2 (pretrained on three species), and a model trained from scratch on the thoracic dataset alone. Global scores are averaged across both muscle classes (meso/metathoracic and prothoracic).

## 4 Discussion

### 4.1 Multi-species generalization within Orthoptera

These results demonstrate that a segmentation model, trained through either a multi-species fine-tuning strategy or simultaneous *from scratch* training, generalizes effectively across several phylogenetically related orthopteran species. Learning through fine-tuning, by incorporating segmentations from different species, sometimes comes with a trade-off between generalization and performance, leading to catastrophic forgetting [8]. Even so, ModelB reaches a mean Dice of 0.7715 across the three validation species combined, without any marked degradation on the reference species even after the progressive integration of *L. equestris* and *L. doenitzi*.

One noteworthy result concerns the InCuticle class, whose Dice coefficient rises from 0.30 (*G. bimaculatus* 01) to 0.54–0.57 as additional individuals are added. This class appears difficult to improve, for a couple of likely reasons. First, the tissue itself is extremely thin, sometimes falling below the size of a single voxel in certain species or specimens, which complicates both manual segmentation and model learning. Second, the internal cuticle is closely associated with the mandibular and antennal muscles, sharing a texture and coloration similar to muscle tissue, which increases the difficulty of producing consistent segmentation across individuals, for both the learning process and the validation tests.

These interpretations should nonetheless be qualified by an alternative explanation: since *Loxoblemmus* specimens were annotated later in the project, their ground truth may have benefited from a potentially different, or even better, annotation method compared to the earliest *G. bimaculatus* specimens. The improvement in Dice for InCuticle could therefore reflect, at least in part, an improvement in the quality of the ground truth rather than genuine model generalization.

The specimen *G. bimaculatus* 02 (female), the second Gryllus validation individual, annotated toward the end of this research, shows a markedly higher InCuticle Dice score (0.53–0.55) than *G. bimaculatus* 01 (0.28–0.30), an interesting observation which, combined with a visual check, confirms that a real change did occur in the manual segmentation method over time, as shown in Figure 9.

**Figure 9:**
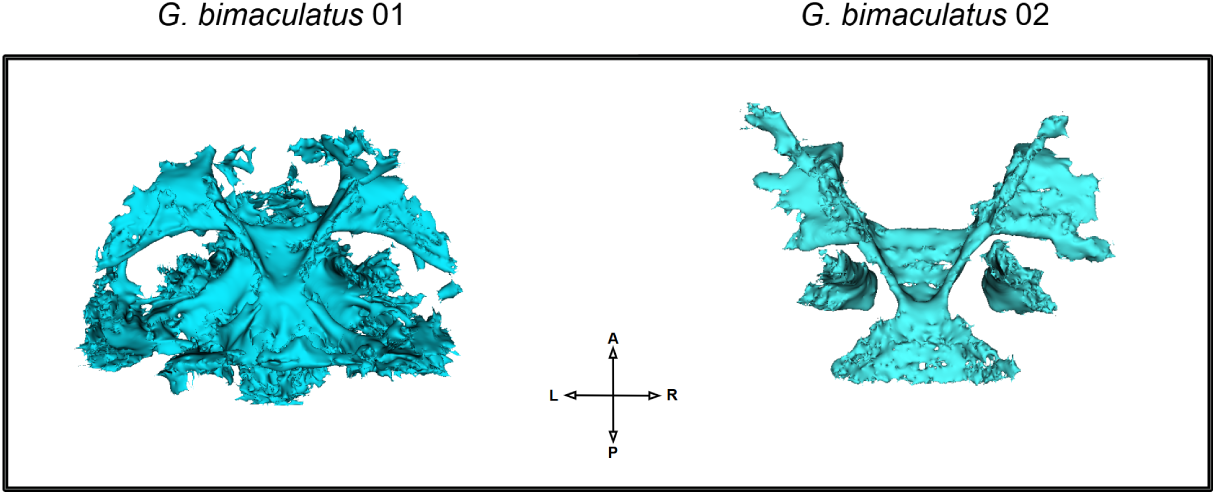
Visual comparison of the InCuticle class between *G. bimaculatus* 01 and *G. bimaculatus* 02.

Unlike this study, where reference segmentation was performed manually for most specimens, Toulkeridou et al. [4] used a standardized semi-automatic segmentation method followed by manual post-processing, which limits variability in annotation quality over time compared to a largely manual approach. This methodological difference highlights the value of semi-automatic approaches (requiring only a minimal amount of segmentation to be performed) for ensuring greater consistency in the ground truth, particularly for thin tissues such as the internal cuticle.

### 4.2 The limits of sequential fine-tuning

Sequential fine-tuning (ModelB) and *from scratch* training (ModelC) produced statistically equivalent performance on the cephalic segmentation task (paired Wilcoxon signed-rank test, n = 24, V = 191.5, p = 0.243). This result indicates that, for this task, neither the order nor the number of previously segmented individuals introduced during training has a significant effect on the model’s final performance, contrary to the initial hypothesis that progressively introducing species would allow the model to better consolidate shared features before incorporating inter-species variability. This result is consistent with the work of González et al. [9], who similarly report a minimal performance difference between a model trained by sequential fine-tuning and one trained *from scratch*, in a medical segmentation context that also uses nnU-Net.

This equivalence did not hold, however, when the approach was extended experimentally to a model trained on the thoracic muscle groups of *G. bimaculatus*. Although both t1 and t2 achieved better results than training from scratch, the model pretrained on two species (t1, Dice = 0.673) outperformed the one pretrained on three species (t2, Dice = 0.633), a counterintuitive result suggesting that the additional diversity introduced by *L. doenitzi* in ModelC’s pretraining did not translate into features more transferable to this specialized thoracic muscle model, and may even have introduced a slightly unfavorable effect. However, the number of individuals manually segmented for this exploratory model was limited to only three, comprising two specimens used for training and a single validation specimen, precluding any formal statistical testing of this difference.

### 4.3 Texture-driven generalization beyond Orthoptera

These results extend and complement those of Toulkeridou et al. [4], who had demonstrated generalization of a 2D U-Net model trained on the ant brain to other nervous structures and other insect orders, without however quantifying this generalization. This work provides a rigorous quantitative evaluation of that same capacity, on a model that simultaneously segments six anatomical structures rather than a single one, and relying on a three dimensional architecture (nnU-Net 3D fullres) rather than a reconstruction built from individual 2D slices, a methodological distinction that allows the network to directly exploit the full spatial context during learning.

The generalization test, carried out without any prior training, on specimens of Hymenoptera (*Camponotus pennsylvanicus*, shown in Figure 7) and Neuroptera (*Coniopteryx pygmaea*) further confirms the observation of Toulkeridou et al. [4] according to which the model’s generalization targets tissue texture rather than anatomical position. Muscles and the external cuticle, structures whose texture is visually shared across insect orders, were recognized, including in the thorax, abdomen, and locomotor limbs, and therefore with no direct equivalent in Orthoptera, whereas positionally specific structures (esophagus, brain, eyes, and internal cuticle) were not correctly localized. This limitation illustrates that the generalization observed within Orthoptera likely rests on a combination of textural similarity and conservation of overall anatomical organization, two factors that dissociate markedly as phylogenetic distance increases at the order level.

### 4.4 Methodological limitations

Several limitations deserve mention. First, the model relies on only 20 individuals for training, validation, and testing combined. Indeed, adding specimens from the same or from another species would increase the model’s accuracy or its generalization. In addition, Toulkeridou et al. [4] use a 60/40 ratio of individuals for learning/validation, compared to 80/20 in this work, which likely constrains our results. Additionally, the statistical power of comparisons between training strategies remains modest: although adding a second *G. bimaculatus* validation specimen allowed the sample to grow from 18 to 24 paired observations, this number remains limited by the usual standards of statistical analysis, and each training method was trained only once. Several random replications per condition would be needed to distinguish a genuine methodological effect from mere stochastic variance between training runs. Third, the InCuticle class remains the hardest to segment across every model tested, with Dice coefficients systematically lower than the other classes despite the progressive improvement of manual segmentation. Even imprecise, the automated segmentation remains visually useful enough for creating templates for manual segmentations.

### 4.5 Perspectives

These results open up several directions for future work. The statistical limitation to be lifted concerns the number of specimens dedicated to validation: annotating four to six additional specimens per species, specifically reserved for validation rather than training, would provide sufficient statistical power for robust comparisons between training strategies and between species, currently limited by a relatively restricted number of test specimens.

On the taxonomic side, extending the model to a larger number of orthopteran species, in particular taxa that are phylogenetically more distant within the order, such as representatives of the suborder Caelifera (grasshoppers), as opposed to the Ensifera (crickets) studied here, would help better characterize the generalization limits identified in this work and evaluate the feasibility of a truly generalist model at the scale of the order Orthoptera. It is still worth noting that the automated segmentation produced by ModelB for a male *L. doenitzi* with pronounced sexual dimorphism (Figures 3 and 4), which had not been seen before, showed surprisingly good predictive power. This assessment remains purely qualitative, however, as no ground truth was available to compute a Dice score.

The semi-automatic workflow developed during this project, combining model prediction with manual correction, is a directly reusable tool for speeding up the annotation of new specimens, significantly reducing manual segmentation time and increasing consistency between individuals, thereby facilitating the future extension of the dataset. Finally, on the applied side, the ability of this kind of tool to rapidly characterize the anatomy of multiple orthopteran species could facilitate the comparative study of their functional traits, their development, their sensory morphology, and other structures related to bioturbation, with a view to better understanding their role in agroecosystems.

## 5 Conclusion

This work developed a 3D, multi-class automated segmentation pipeline for the orthopteran head in micro-CT, based on the nnU-Net framework, and evaluated its ability to generalize across three phylogenetically related species (*Gryllus bimaculatus*, *Loxoblemmus equestris*, and *L. doenitzi*). The final model reaches an overall Dice coefficient of 0.7715 across the three combined species.

Comparing two training strategies, sequential fine-tuning by species (ModelB) and simultaneous *from scratch* training (ModelC), revealed no statistically significant difference for head tissue segmentation. An opposite trend was nonetheless observed on an anatomically distinct task, thoracic muscle segmentation, where the model pretrained on a smaller number of species (t1) outperformed the one pretrained on a larger number of species (t2). This contrast suggests that the benefit of multi-species transfer likely depends on how closely related the source pretraining task (cephalic segmentation) is to the target adaptation task (thoracic segmentation).

Generalization tests conducted beyond Orthoptera further showed that the model’s ability to recognize structures in more distantly related taxa relies mainly on tissue texture rather than anatomical position, confirming observations previously reported in the literature for other insect groups.

This work therefore demonstrates the feasibility of a generalizable automated segmentation approach within a single insect order, while also identifying part of the limits of that generalization, and provides a methodological foundation that can be directly reused to extend the model to other orthopteran species and to accelerate the annotation of new datasets in comparative morphology.

## Data Availability

The trained models are available at Zenodo: https://doi.org/10.5281/zenodo.21617087. The source code is also maintained on GitHub: https://github.com/arthurcheron/orthoptera-head-segmentation.

## Acknowledgments

We thank Drs. Jonchee Kao and Takaaki Daimon, and everyone in the laboratory for their assistance and discussion. We would also like to thank PhD candidate Tancrède Martinez Boisseau for all the answers provided regarding deep learning and the technical issues encountered. This work was supported by the M1 AETPF internship program of Université Paris-Saclay and JST PRESTO Grant Number JPMJPR25N5 and NIBB Collaborative Research Program Grant Numbers 25NIBB303 and 26NIBB305.

## A Appendices

### A.1 Key Python scripts

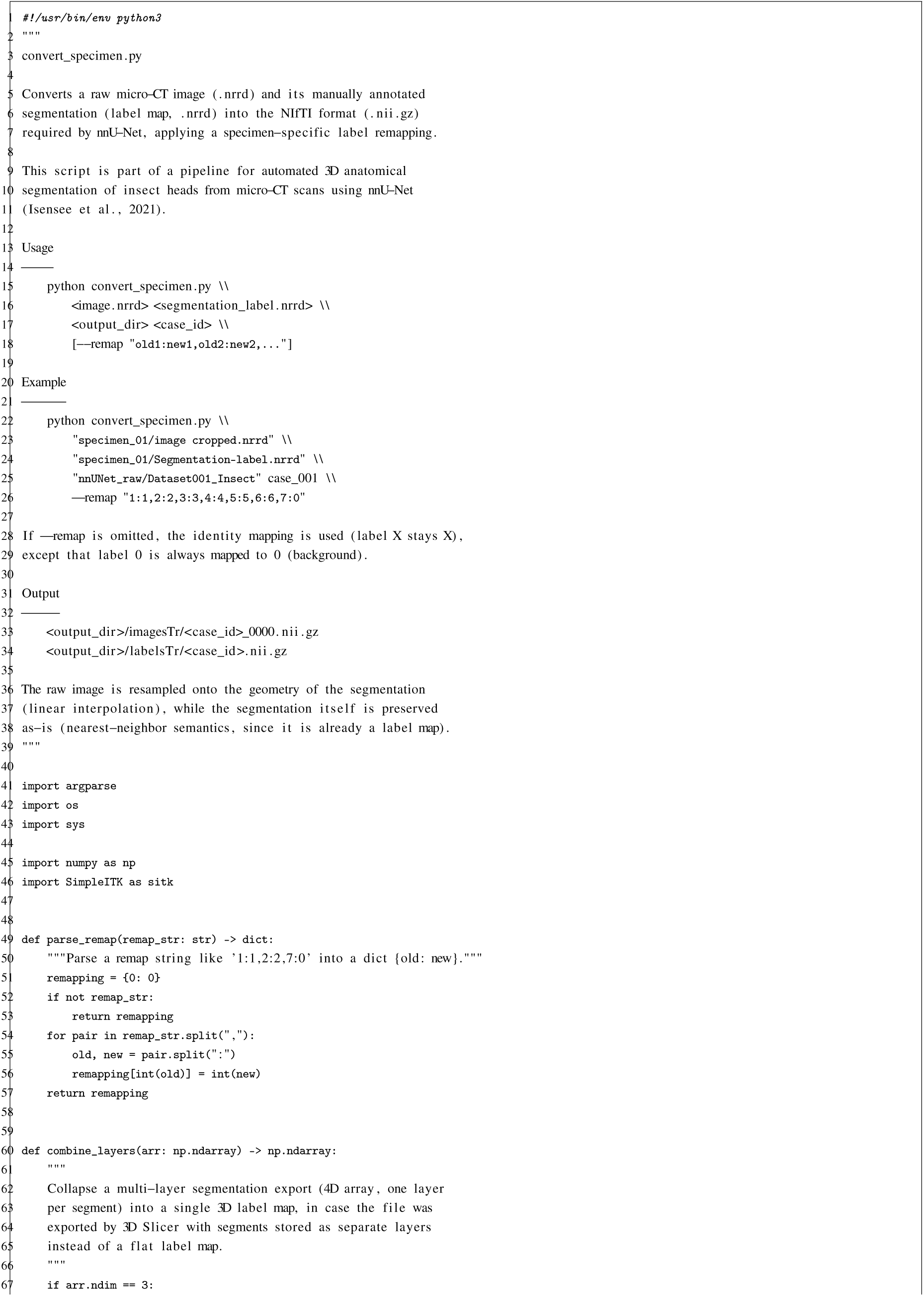
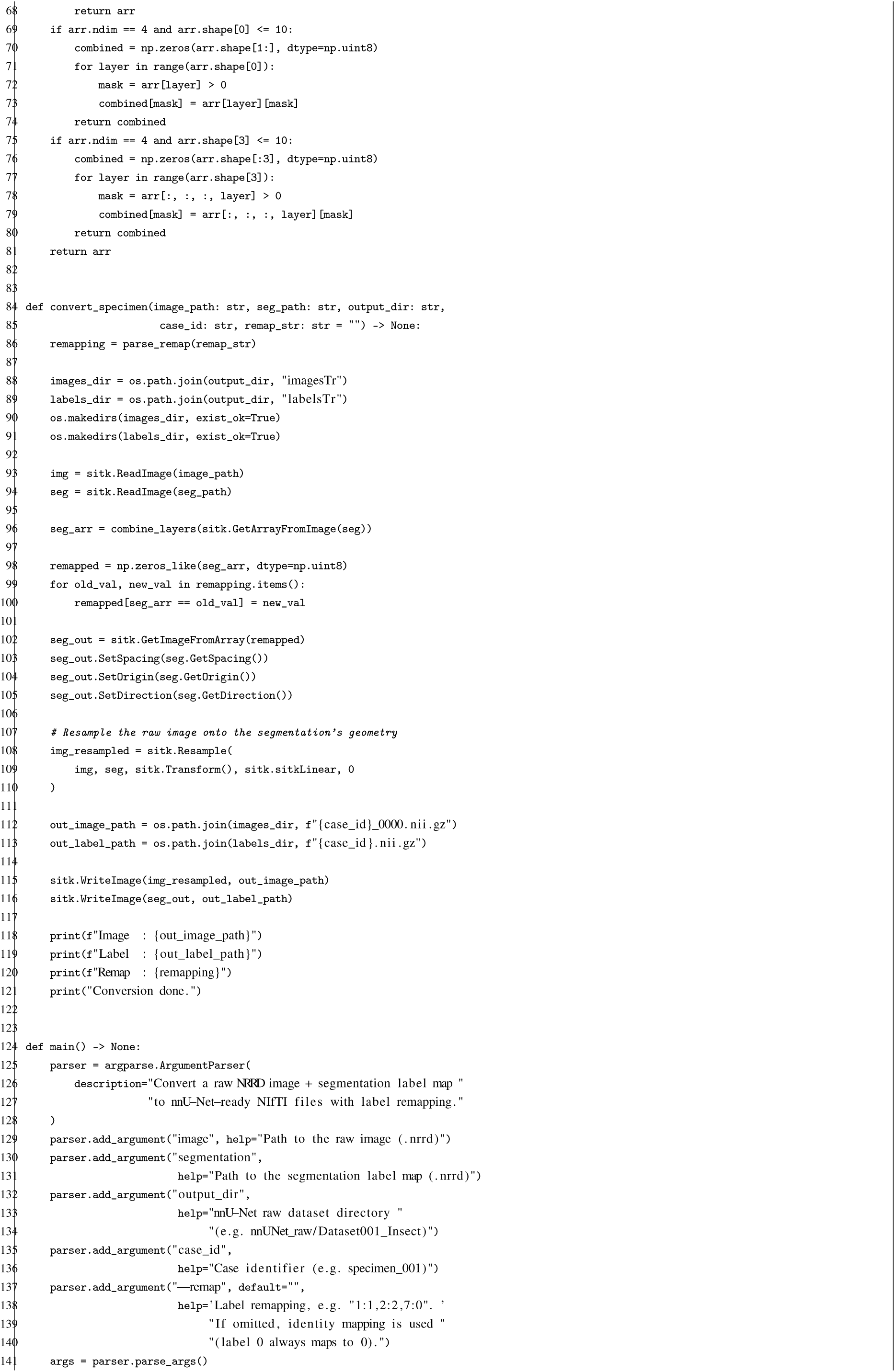
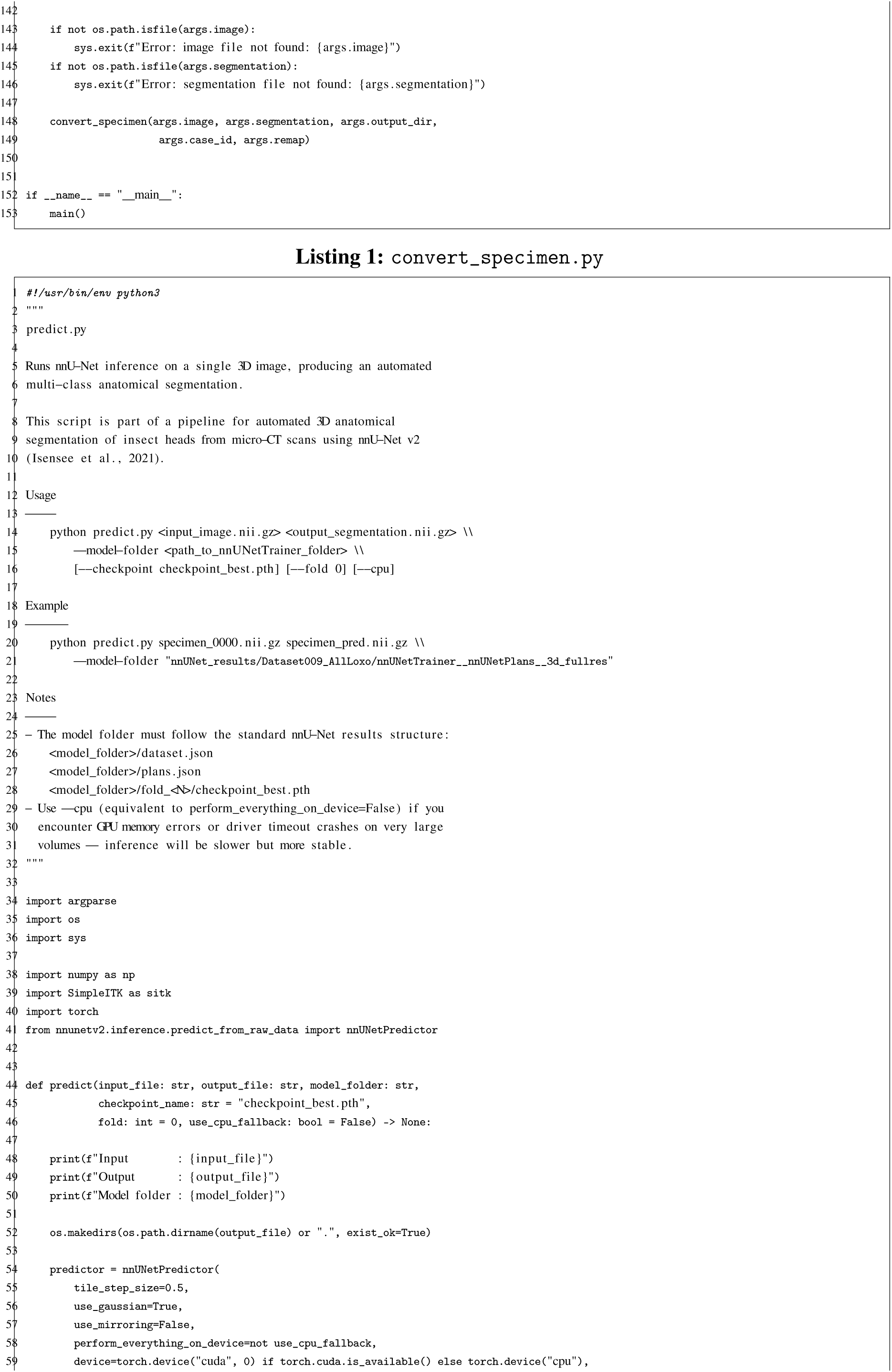
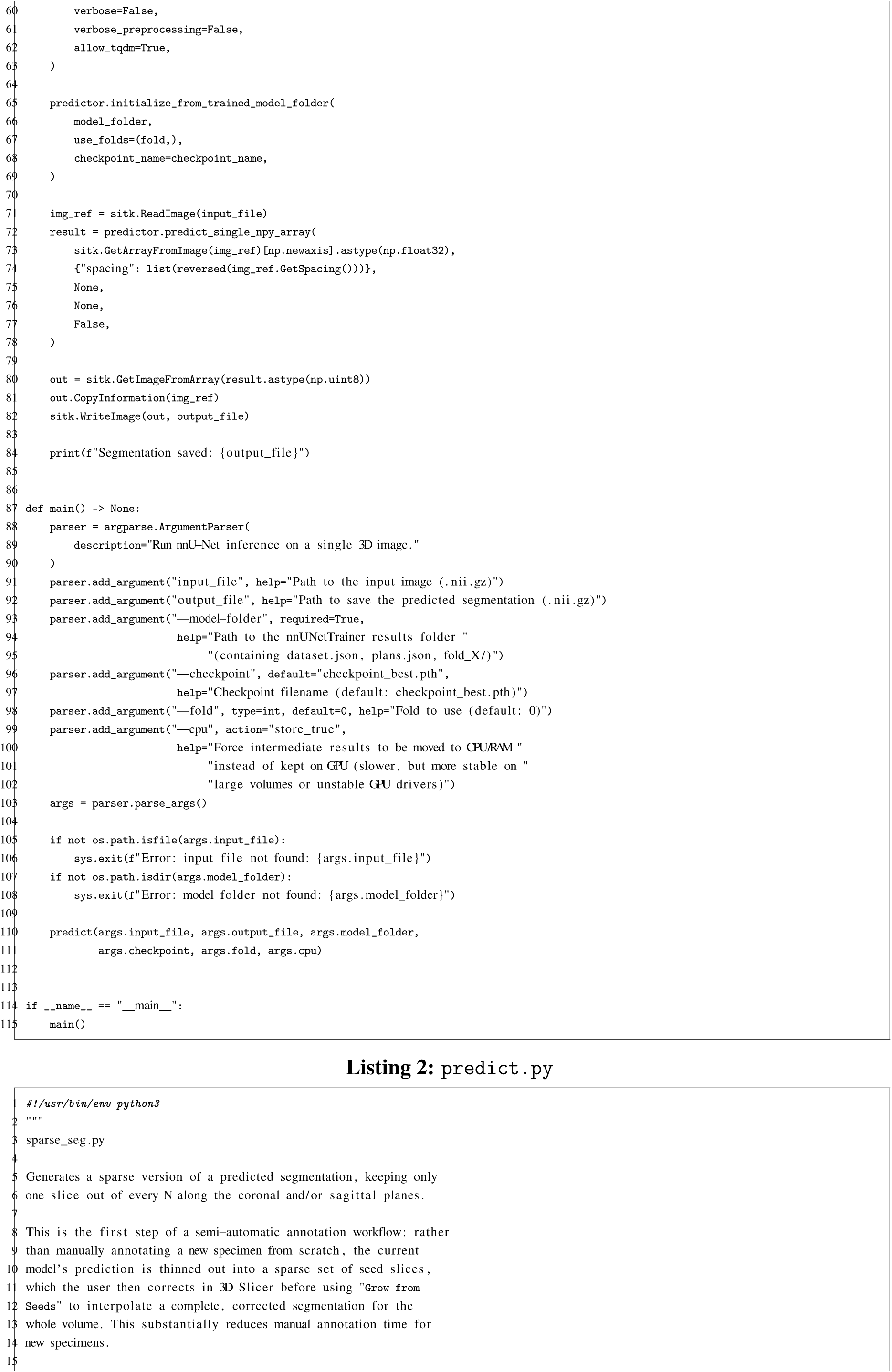
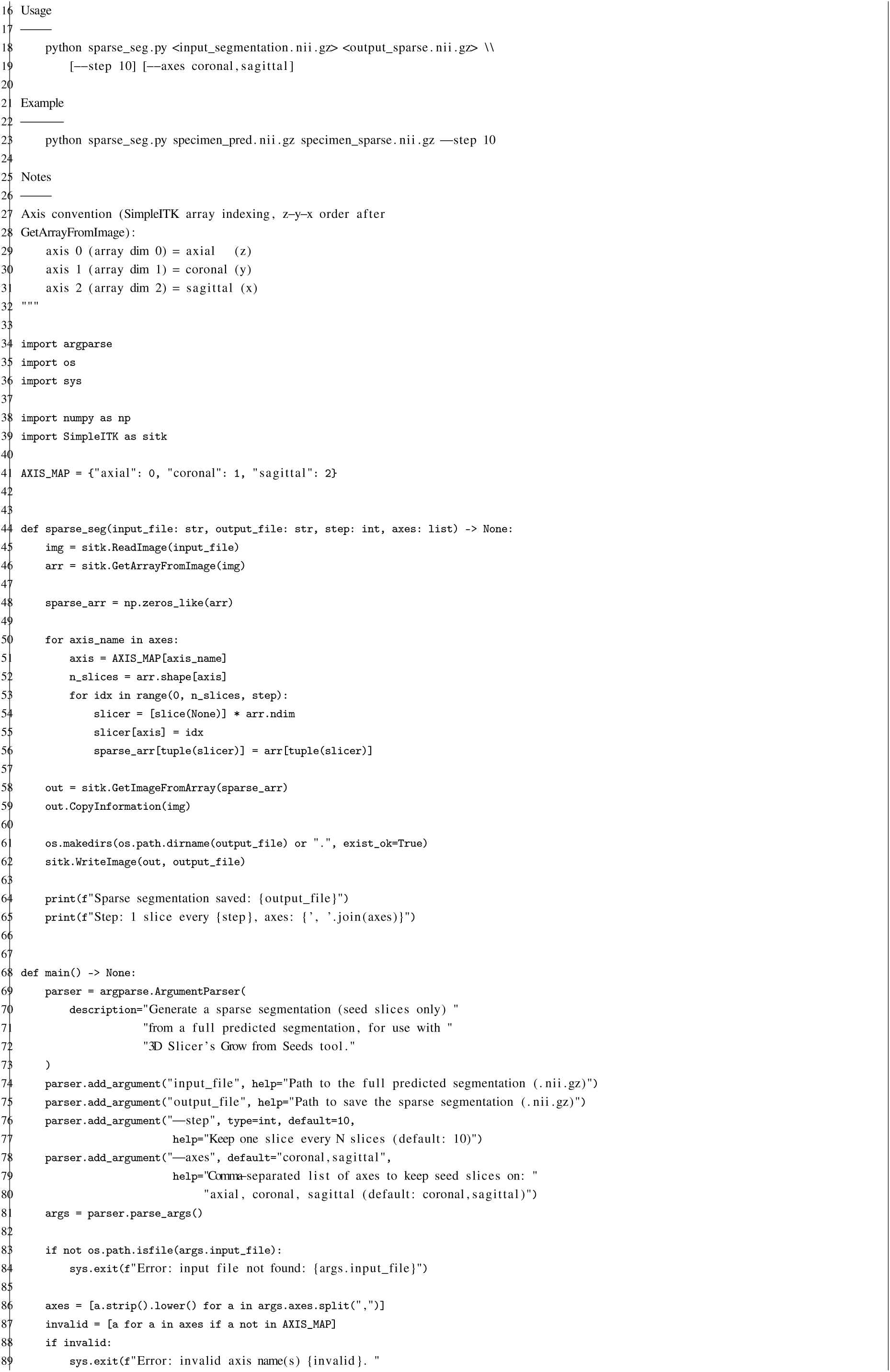
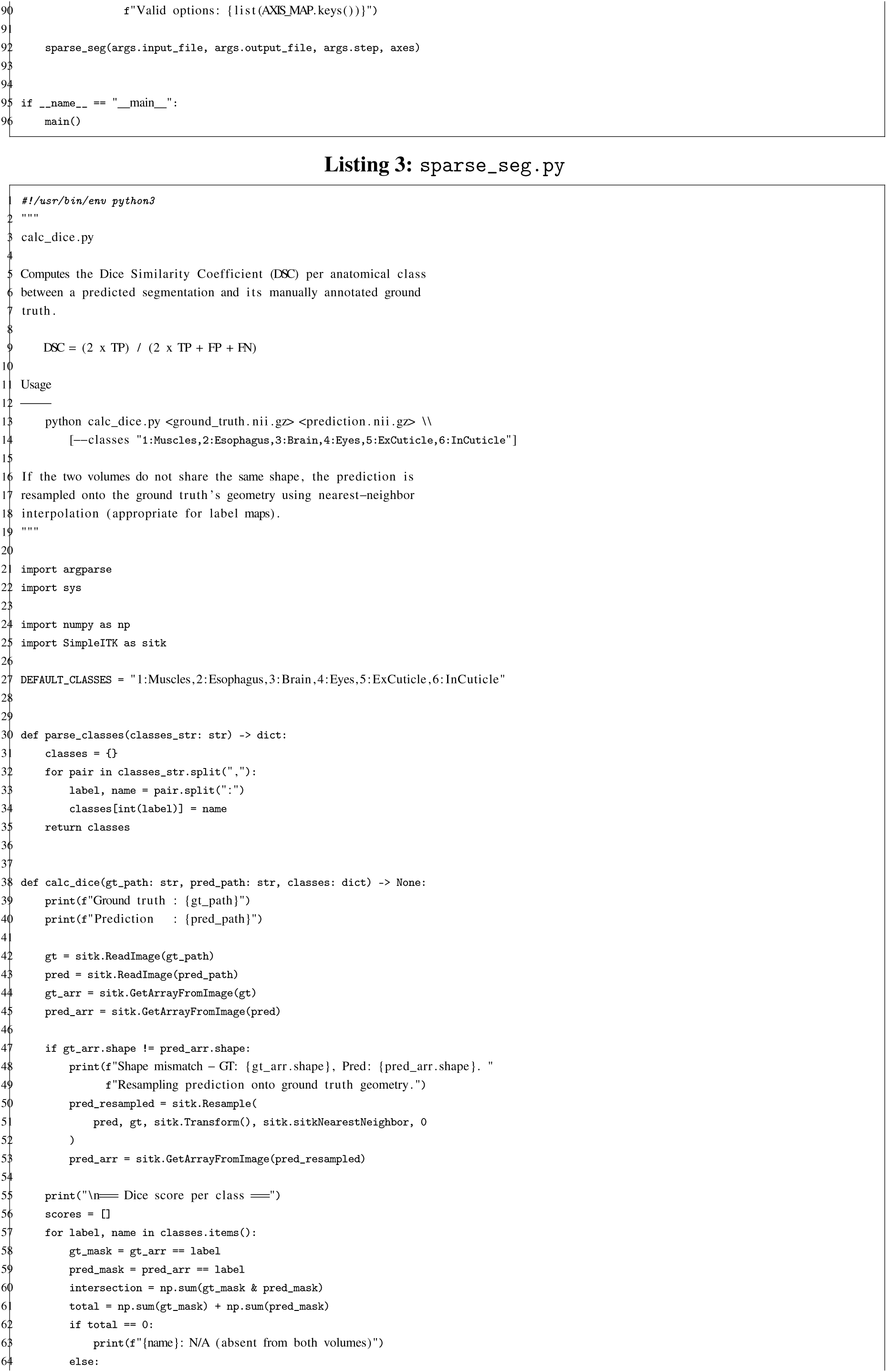
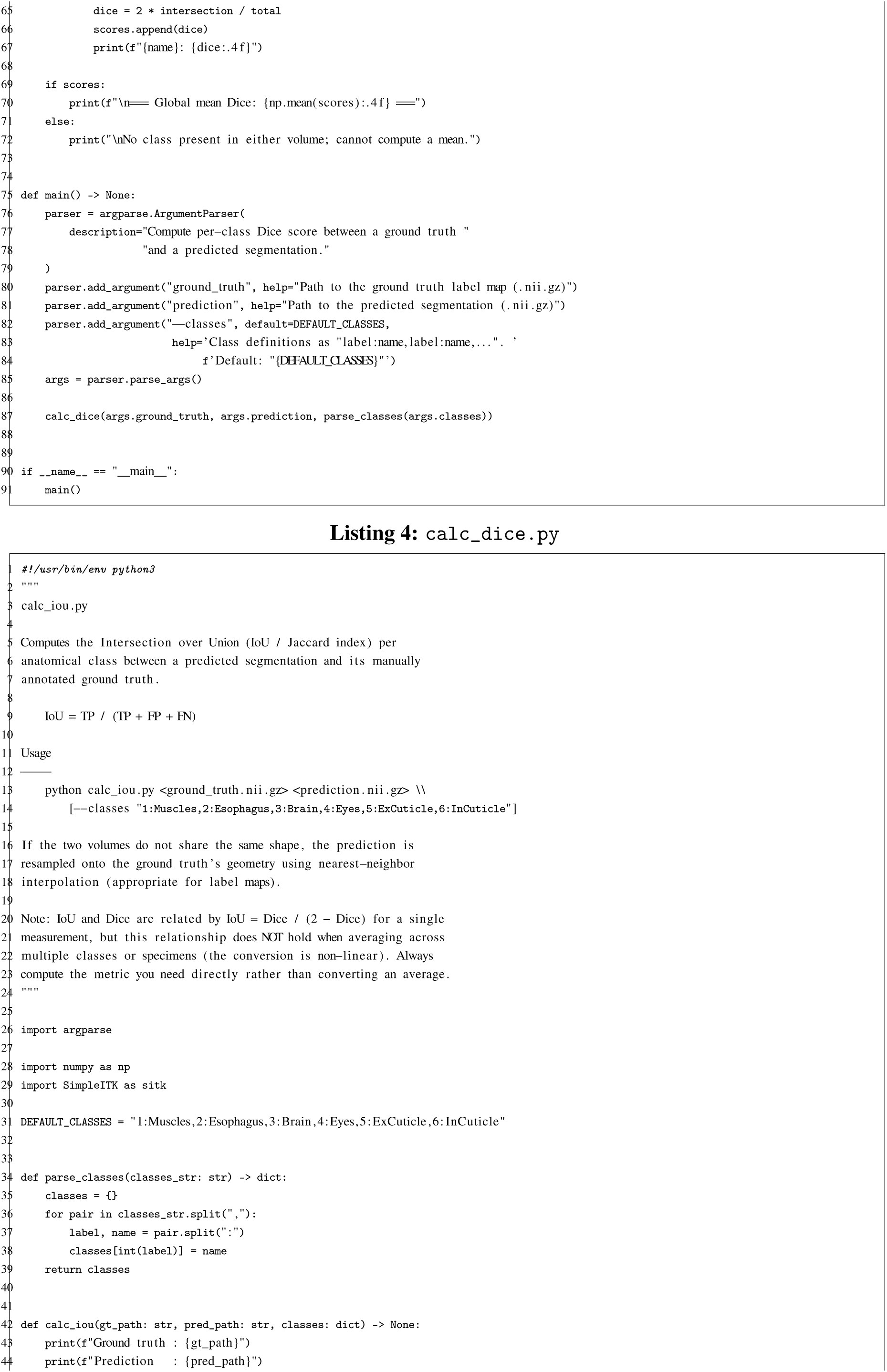
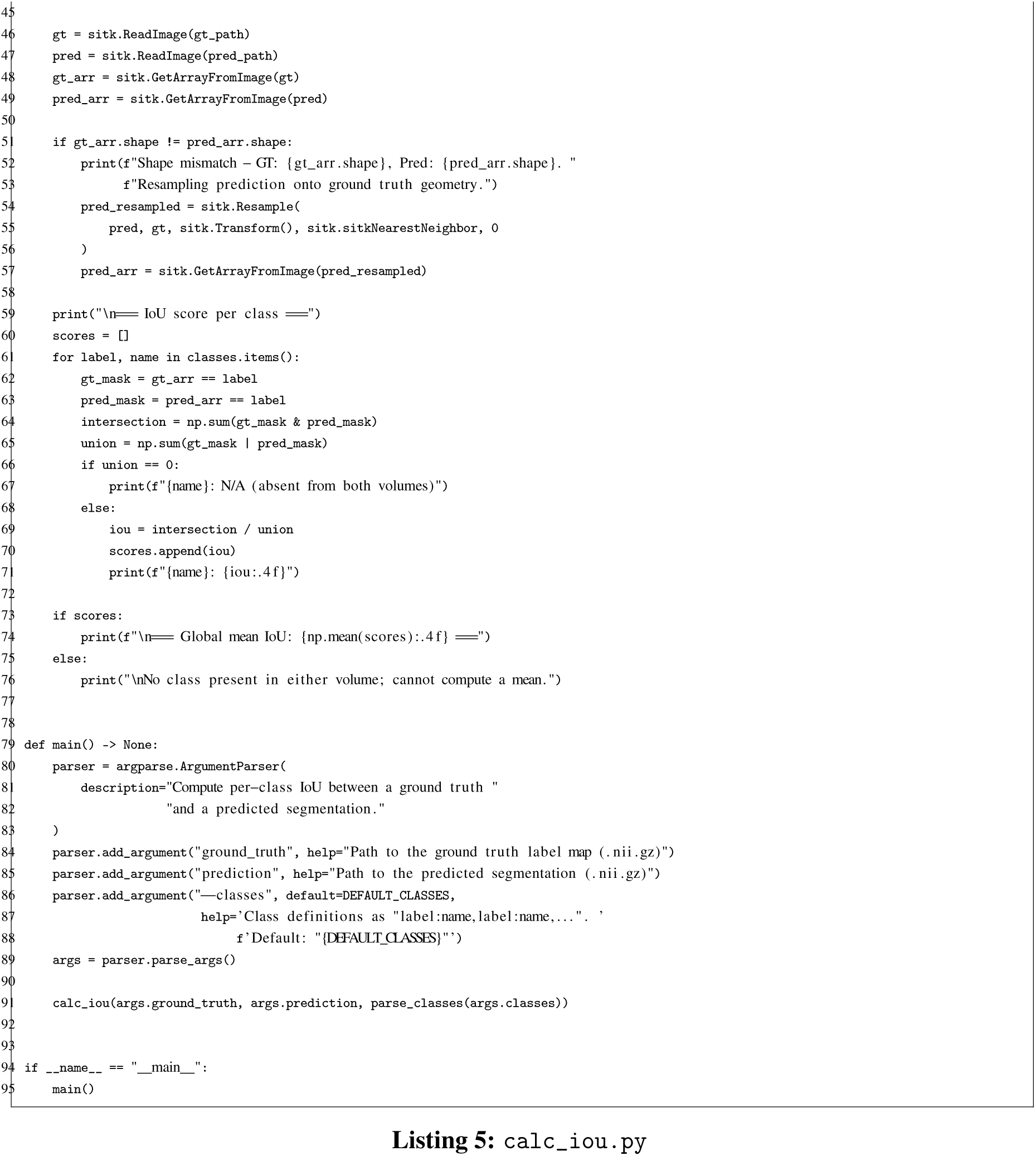

### A.2 Complete tracking tables

**Table 1:** Complete list of specimens used for training, validation, and testing across the three species studied, plus one *Camponotus pennsylvanicus* specimen used only for the cross-order generalization test (Figure 7). “Voxel size” is the spacing measured in the NIfTI files used for training (after cropping/exporting from 3D Slicer); “Native voxel size” was verified directly in 3D Slicer on the raw files. For cricket_012, the recorded native voxel size (0.198 mm) is inconsistent with the expected physical dimensions of an insect head and remains to be clarified; this specimen is excluded from both training and validation.

| Internal ID | Species | Stage | Voxel size (mm) | Native voxel size (mm) | Volume dimensions | Role (Model-B) | Role (Model-A) | Notes |
| --- | --- | --- | --- | --- | --- | --- | --- | --- |
| cricket_001 | *Gryllus bimaculatus* | Adult male | 0.022584 | 0.005646 | 278x312x257 | Training | Training | – |
| cricket_002 | *Gryllus bimaculatus* | Adult male | 0.005615 | 0.005615 | 641x596x542 | Training | Training | – |
| cricket_003 | *Gryllus bimaculatus* | Adult female | 0.008003 | 0.008003 | 1008x1008x630 | Training | Training | – |
| cricket_004 | *Gryllus bimaculatus* | Adult female | 0.008 | 0.008 | 871x759x650 | Validation | Validation | Validation specimen kept constant across all model versions |
| cricket_005 | *Gryllus bimaculatus* | Adult female | 0.008 | 0.008 | 822x772x603 | Training | Training | – |
| cricket_006 | *Gryllus bimaculatus* | Adult female | 0.008 | 0.008 | 822x772x604 | Excluded | Excluded | Duplicate scan of cricket_005 |
| cricket_007 | *Gryllus bimaculatus* | Adult male | 0.022584 | 0.006561 | 282x320x233 | Training | Training | – |
| cricket_008 | *Gryllus bimaculatus* | Adult male | 0.02246 | 0.02246 | 160x151x138 | Training | Training | Highest resolution |
| gryllus_female02 | *Gryllus bimaculatus* | Adult female | 0.017996 | 0.017996 | 345x397x296 | Held-out test | – | – |
| cricket_009 | *Loxoblemmus equestris* | Adult male | 0.011744 | 0.006 | 312x274x220 | Training | Training | – |
| cricket_010 | *Loxoblemmus equestris* | N9 nymph | 0.007002 | 0.001751 | 496x397x536 | Training | Training | – |
| cricket_011 | *Loxoblemmus equestris* | N9 nymph | 0.006101 | 0.006101 | 718x584x421 | Training | Training | – |
| cricket_012 | *Loxoblemmus equestris* | Adult male | 0.002 | 0.198 | 404x467x296 | Excluded | Excluded | Domain shift (atypical contrast) |
| cricket_013 | *Loxoblemmus equestris* | N9 nymph | 0.006401 | 0.0016 | 423x523x550 | Training | Training | – |
| cricket_014 | *Loxoblemmus equestris* | Adult female | 0.0054 | 0.00135 | 651x436x651 | Training | Training | – |
| cricket_015 | *Loxoblemmus equestris* | N9 nymph, female | 0.006002 | 0.001501 | 547x503x431 | Validation | Training | – |
| doenitzi_001 | *Loxoblemmus doenitzi* | Last instar nymph, female | 0.0064 | 0.0016 | 811x797x495 | Training | – | Annotated via assisted workflow (sparse + Grow from Seeds) |
| doenitzi_002 | *Loxoblemmus doenitzi* | Last instar nymph, male | 0.0064 | 0.0016 | 774x915x529 | Training | – | Annotated via assisted workflow (sparse + Grow from Seeds) |
| doenitzi_003 | *Loxoblemmus doenitzi* | Last instar nymph, male | 0.007 | 0.007 | 734x801x559 | Validation | – | Complete manual annotation, dedicated to validation |
| doenitzi_004 | *Loxoblemmus doenitzi* | Adult male | 0.0096 | 0.0024 | 803x870x408 | Training | – | Annotated via assisted workflow (sparse + Grow from Seeds) |
| doenitzi_005 | *Loxoblemmus doenitzi* | Adult male | 0.0096 | 0.0024 | 868x835x370 | Training | – | Annotated via assisted workflow (sparse + Grow from Seeds) |
| doenitzi_006 | *Loxoblemmus doenitzi* | Adult male | 0.019824 | 0.004956 | 425x432x228 | Training | – | Annotated via assisted workflow (sparse + Grow from Seeds) |
| Camponotus | *Camponotus pennsylvanicus* | Worker | 0.005516 | 0.005516 | 1495x1012x929 | Visualization test | – | Predict by ModelB for visualization test |

